# Antimycin A induces light hypersensitivity of photosystem II in the presence of Q_B_-site binding herbicides

**DOI:** 10.1101/2024.08.26.609723

**Authors:** Ko Imaizumi, Daisuke Takagi, Kentaro Ifuku

## Abstract

Photosynthetic electron transport consists of linear electron flow and two cyclic electron flow (CEF) pathways around photosystem I (PSI). PGR5-dependent CEF-PSI is thought to be the major CEF-PSI pathway and an important regulator of photosynthetic electron transfer. Antimycin A (AA) is commonly recognized as an inhibitor of PGR5-dependent CEF-PSI in photosynthesis. Although previous findings imply that AA may also affect photosystem II (PSII), which does not participate in CEF-PSI, these “secondary effects” tend to be neglected, and AA is often used for inhibition of PGR5-dependent CEF-PSI as if it were a specific inhibitor. Here, we investigated the direct effects of AA on PSII using isolated spinach PSII membranes, and spinach and *Chlamydomonas* thylakoid membranes. Measurements of Q_A_^−^ reoxidation kinetics showed that AA affects the acceptor side of PSII and inhibits electron transport within PSII. Furthermore, repetitive *F*_v_/*F*_m_ measurements revealed that, in the presence of Q_B_-site binding inhibitors, AA treatment results in severe photoinhibition even from a single-turnover flash. The direct effects of AA on PSII are non-negligible and caution is required when using AA as an inhibitor of PGR5-dependent CEF-PSI. Meanwhile, we found that the commercially available compound AA3, which is a component of the AA complex, inhibits PGR5-dependent CEF-PSI without having notable effects on PSII. Thus, we propose that AA3 should be used for the physiological study of the PGR5-dependent CEF-PSI.

## Introduction

In oxygenic photosynthesis, photosystem II (PSII), a pigment–protein complex, uses light energy to oxidize water to molecular oxygen at its donor side, and reduce plastoquinone (PQ) molecules at its acceptor side (Shen, 2015). Upon absorption of light energy, excitation of P680, the reaction center chlorophylls of PSII, is followed by charge separation (Yoneda *et al*., 2022; Nguyen *et al*., 2023). The generated highly oxidative electron hole drives water oxidation at the Mn_4_CaO_5_ cluster, while the generated electron is transferred from pheophytin (Pheo_D1_) to the non-exchangeable primary electron acceptor quinone Q_A_, and further to the exchangeable secondary electron acceptor quinone Q_B_ (Shevela *et al*., 2023) (**Fig. 1**). Upon acceptance of a second electron, Q_B_^−^ takes up two protons and dissociates from the Q_B_-site of PSII as plastoquinol (PQH_2_), thus entering the PQ pool.

**Fig. 1.**
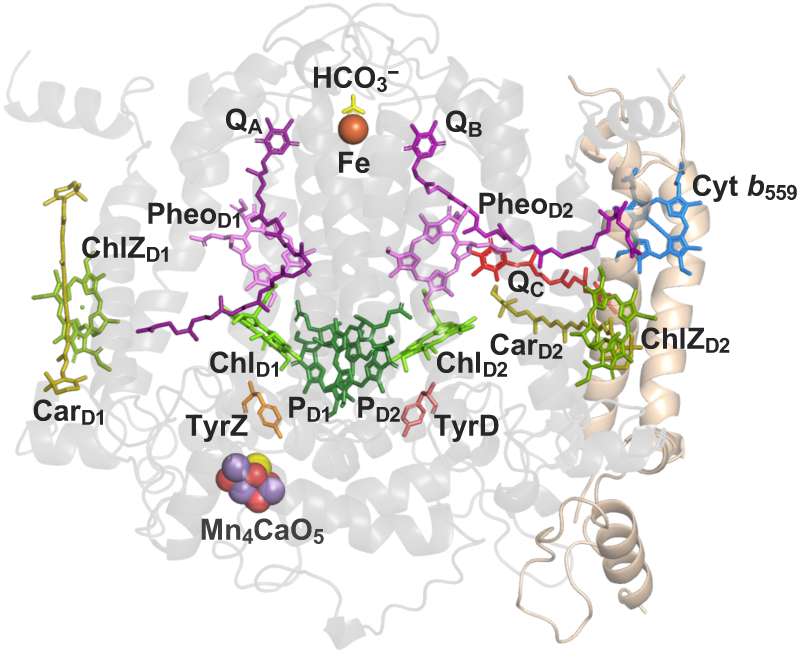
Electron transfer cofactors in PSII. The electron transfer cofactors, other redox-active cofactors, and related components in the reaction center of a PSII monomer are shown. P680 (reaction center chlorophylls P_D1_, P_D2_, Chl_D1_ and Chl_D2_), pheophytins (Pheo_D1_ and Pheo_D2_), plastoquinones (Q_A_ and Q_B_), redox-active tyrosines (TyrZ and TyrD), extra chlorophylls (ChlZ_D1_ and ChlZ_D2_), associated carotenoids (Car_D1_ and Car_D2_), and the bicarbonate ion are shown as stick models. The Mn_4_CaO_5_ cluster and the non-heme iron are shown as spheres. In addition, the heme of Cyt *b*_559_ and the third plastoquinone molecule (Q_C_) located in a quinone exchange channel are shown as stick models. The D1 and D2 subunits are shown in gray and the Cyt *b*_559_ α- and β-subunits (PsbE and PsbF) are shown in beige cartoon view. The PSII structure was generated using PDB ID: 3WU2, with only the Q_C_ molecule generated from PDB ID: 4V62.

In the dominant photosynthetic electron transport pathway, linear electron flow, electrons pass sequentially from PQH_2_ through cytochrome (Cyt) *b*_6_*f* and plastocyanin to PSI, and further to ferredoxin (Fd), and are used for reduction of NADP^+^ to NADPH by Fd-NADPH^+^ oxidoreductase (FNR). Cyclic electron flow around PSI (CEF-PSI) is an important alternative electron transport pathway (Yamori and Shikanai, 2016). In CEF-PSI, PSI reduces Fd and electrons are transported from Fd back to the PQ pool. Of several proposed CEF-PSI pathways, two well-known pathways are the NDH-dependent pathway (Burrows *et al*., 1998; Kofer *et al*., 1998; Shikanai *et al*., 1998) and the PGR5-dependent pathway (or PGR5/PGRL1-dependent pathway) (Munekage *et al*., 2002, 2004; DalCorso *et al*., 2008). The NDH-dependent pathway is mediated by the NAD(P)H dehydrogenase-like (NDH) complex (Ifuku *et al*., 2011; Peltier, Aro and Shikanai, 2016), whereas the PGR5-dependent pathway requires the protein PROTON GRADIENT REGULATION 5 (PGR5) (Munekage *et al*., 2002). Recent progress has been achieved in understanding the mechanisms of the NDH-dependent CEF-PSI (Shen *et al*., 2022; Zhang *et al*., 2023). In contrast, although the importance of PGR5 has been revealed (Yamori and Shikanai, 2016; Ma *et al*., 2021), the molecular mechanism of the PGR5-dependent CEF-PSI pathway remains elusive, leading to controversy over its exact role (Nandha *et al*., 2007; Suorsa *et al*., 2012; Leister and Shikanai, 2013; Fisher and Kramer, 2014; Takagi and Miyake, 2018; Nawrocki *et al*., 2019; Buchert *et al*., 2020; Rantala *et al*., 2020; Rühle *et al*., 2021; Wada, Amako and Miyake, 2021; Wu *et al*., 2021; Zhou *et al*., 2022).

In the field of photosynthesis, antimycin A (AA) has long been recognized as an inhibitor of CEF-PSI (Tagawa, Tsujimoto and Arnon, 1963). AA has been suggested to inhibit PGR5-dependent CEF-PSI but not NDH-dependent CEF-PSI (Munekage *et al*., 2002, 2004). Subsequently, AA has often been used as an inhibitor of PGR5-dependent CEF-PSI, although neither the target nor the inhibitory mechanism of AA are known. We recently provided evidence that AA lowers the redox potential of Cyt *b*_559_, a component of the PSII core, and affects electron transfer within PSII (Takagi *et al*., 2019), which is considered not to be involved in CEF-PSI. Several previous studies support that AA may influence PSII (Katoh, 1972; Yerkes and Crofts, 1992), especially the properties of Cyt *b*_559_ (Hind, 1968; Cramer, Fan and Böhme, 1971; Cramer and Böhme, 1972; Satoh and Katoh, 1972; Miyake, Schreiber and Asada, 1995). However, these findings have received little attention (Labs, Rühle and Leister, 2016), perhaps due to limited information clarifying the importance of the effects of AA on PSII, as well as due to the lack of alternative inhibitors known to inhibit PGR5-dependent CEF-PSI with a higher specificity. AA remains to be commonly used for inhibiting PGR5-dependent CEF-PSI without sufficient consideration on its possible effects on PSII. Here, we investigated the direct effects of AA on PSII to clarify whether these effects really are negligible, using isolated spinach PSII membranes and spinach and *Chlamydomonas* thylakoid membranes. Furthermore, we propose the use of an alternative inhibitor of PGR5-dependent CEF-PSI, AA3, which we found to have no notable effect on PSII.

## Results

### AA suppresses electron transfer within PSII

We first examined AA effects on the Q_A_^−^ reoxidation kinetics of PSII in isolated spinach thylakoid membranes. The reduction of Q_A_ by a single-turnover flash leads to an increase in fluorescence yield. The subsequent fluorescence decay in the dark, reflecting Q_A_^−^ reoxidation, was monitored to obtain information on electron transfer within PSII. The fluorescence decay after a single-turnover flash can be fitted to three exponential components and a residual, time-independent component (usually with a lifetime > 10 s) (Reifarth *et al*., 1997). In the absence of DCMU, the fastest decay component reflects electron transfer from Q_A_^−^ to Q_B_, the intermediate decay component represents electron transfer from Q_A_^−^ to Q_B_ after a PQ molecule binds to an empty Q_B_-site, the slow decay component reflects Q_A_^−^ charge recombination with the donor side of PSII, and the residual component is attributable to the equilibrium between Q_A_^−^ and Q_B_ (Robinson and Crofts, 1983; Reifarth *et al*., 1997). Because DCMU, AA, and other inhibitor compounds used in this study were dissolved in ethanol, we confirmed that the final concentration of 0.2% (v/v) ethanol did not affect the Q_A_^−^ reoxidation kinetics (**Extended Data Fig. 1**). In the absence of DCMU, AA significantly slowed Q_A_^−^ reoxidation in spinach thylakoid membranes (**Fig. 2a** and **Table 1**). The proportion of the fast phase was strongly reduced, and the time constants for the fast and intermediate phases were more than doubled, implying that AA suppresses electron transfer from Q_A_^−^ to Q_B_ and the binding of PQ to an empty Q_B_-site. In the presence of DCMU, AA had little effect on the proportions of the fast, intermediate, and slow phases, but strongly increased the residual fluorescence level. Similar results were observed using thylakoid membranes isolated from the green alga *Chlamydomonas reinhardtii*, suggesting that the AA effects on PSII are common, at least among green plants (**Extended Data Fig. 2**). Similar effects of AA on PSII were observed with isolated spinach PSII membranes (**Fig. 2b** and **Table 1**), indicating that AA directly binds to PSII and significantly affects electron transfer within PSII. Overall, Q_A_^−^ reoxidation was slightly slower in isolated PSII membranes compared with that in thylakoid membranes, consistent with a previous report, likely due to a higher proportion of PSII having an empty Q_B_-site and presence of fewer unbound PQ molecules in isolated PSII membranes (Roose, Frankel and Bricker, 2010).

**Fig. 2.**
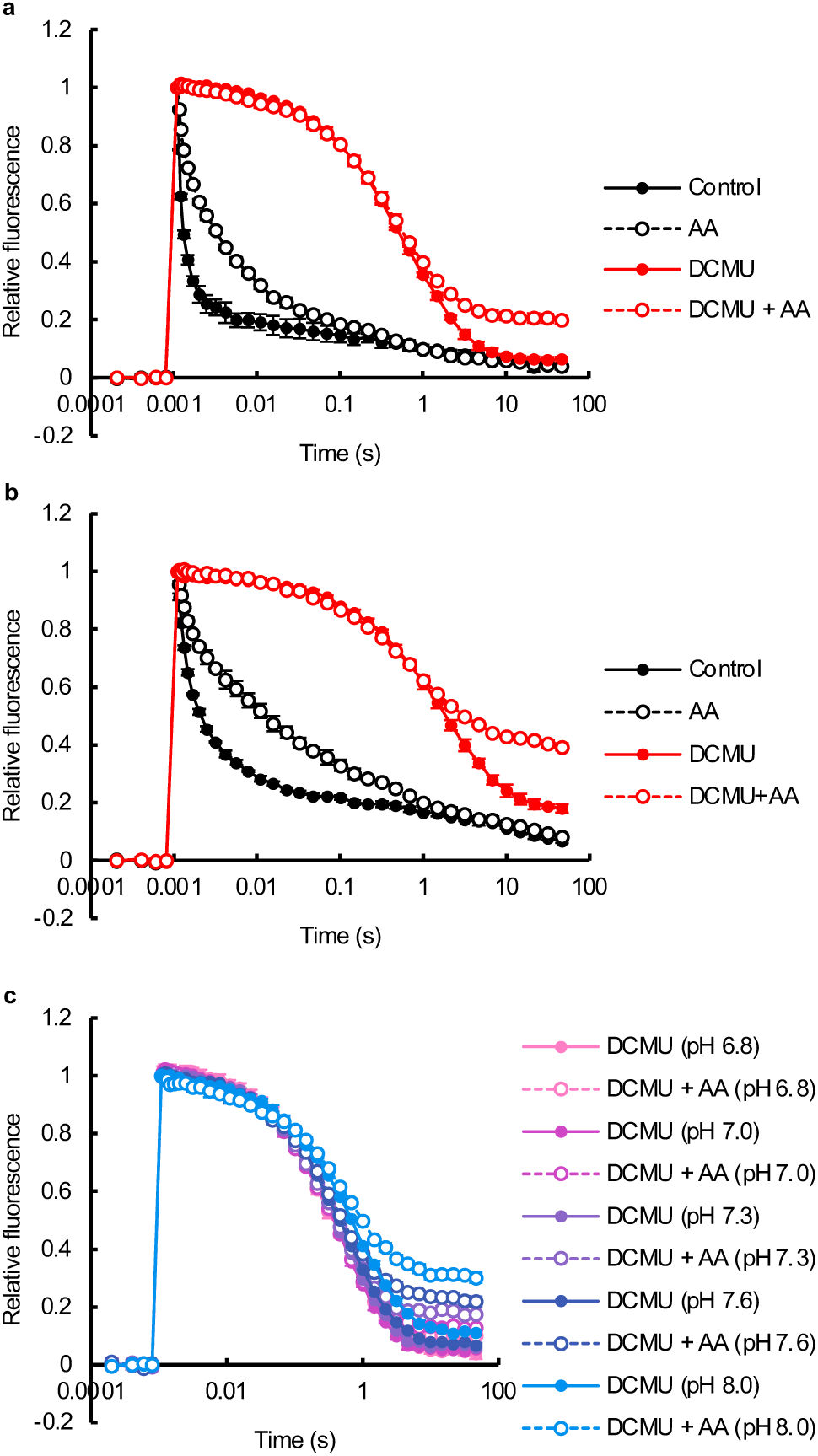
Effects of antimycin A (AA) on the Q_A_^−^ reoxidation kinetics of PSII. (**a**, **b**) Effects of AA on PSII in (**a**) spinach thylakoid membranes and (**b**) isolated spinach PSII membranes in the absence or presence of DCMU. Control (black filled circles), 10 µM AA (black open circles), 10 µM DCMU (red open circles), 10 µM DCMU + 10 µM AA (red open circles). Data are mean ± SD (*n* = 3, independent experiments). (**c**) pH dependence of the effects of AA on the Q_A_^−^ reoxidation kinetics of PSII in spinach thylakoid membranes in the presence of DCMU. Measurements were conducted at pH 6.8 (pink), pH 7.0 (magenta), pH 7.3 (purple), pH 7.6 (blue), pH 8.0 (light blue) in the presence of 10 µM DCMU (filled circles) or 10 µM DCMU + 10 µM AA (open circles). Data are mean ± SD (*n* = 3, technical replicates). Similar results were confirmed in independent experiments. In all panels, in some instances, error bars are smaller than the symbol.

**Table 1.** Kinetic parameters for the Q_A_^−^ reoxidation after a single flash in thylakoid membranes and PSII membranes with or without DCMU and/or antimycin A (AA) treatment.

|  | Fast phase |  | Intermediate phase |  | Slow phase |  | Residual |
| --- | --- | --- | --- | --- | --- | --- | --- |
| | A1 (%) | $\tau_1$ (ms) | A2 (%) | $\tau_2$ (ms) | A3 (%) | $\tau_3$ (s) | y0 (%) |
| <u>Thylakoid</u> |  |  |  |  |  |  |  |
| Control | 77.3 ± 1.3 | 0.15 ± 0.01 | 12.3 ± 0.5 | 2.37 ± 0.09 | 6.9 ± 1.1 | 1.15 ± 0.40 | 3.5 ± 0.6 |
| AA | 43.8 ± 0.4*** | 0.34 ± 0.01*** | 35.7 ± 0.4*** | 6.41 ± 0.68** | 15.3 ± 0.1** | 0.56 ± 0.09 | 5.2 ± 0.5* |
| DCMU | 7.2 ± 0.7 | 27.3 ± 7.1 | 42.0 ± 3.6 | 334 ± 58 | 44.2 ± 4.8 | 1.95 ± 0.26 | 6.6 ± 0.8 |
| DCMU+AA | 6.6 ± 0.9 | 10.7 ± 9.1 | 40.8 ± 3.2 | 304 ± 69 | 33.9 ± 0.8 | 1.55 ± 0.20 | 21.0 ± 0.6*** |
| <u>PSII</u> |  |  |  |  |  |  |  |
| Control | 57.4 ± 0.2 | 0.37 ± 0.02 | 23.8 ± 0.3 | 7.1 ± 0.7 | 11.2 ± 0.4 | 4.0 ± 1.1 | 7.6 ± 0.2 |
| AA | 35.2 ± 2.5** | 0.68 ± 0.10* | 32.2 ± 1.2** | 19.5 ± 1.0*** | 21.1 ± 1.2** | 1.1 ± 0.2* | 11.6 ± 0.5** |
| DCMU | 7.9 ± 1.6 | 63.3 ± 16.5 | 37.0 ± 1.7 | 1016 ± 190 | 36.2 ± 4.2 | 5.4 ± 0.8 | 18.8 ± 1.2 |
| DCMU+AA | 8.0 ± 1.5 | 27.5 ± 9.1* | 32.3 ± 5.4 | 628 ± 234 | 19.2 ± 5.6* | 5.4 ± 3.2 | 40.5 ± 1.4*** |

(A1, $\tau_1$ ), (A2, $\tau_2$ ), and (A3, $\tau_3$ ) are (% amplitudes, exponential decay time constants) of the fast, intermediate, and slow phases; y0 is the % amplitude of the residual fraction. Asterisks indicate statistically significant differences after the addition of AA (Welch's t test, \* $p < 0.05$ , \*\* $p < 0.01$ , \*\*\* $p < 0.001$ , $n = 3$ , independent experiments).

While the observed effects of AA on the Q_A_^−^ reoxidation kinetics with thylakoid membranes in the absence of DCMU were consistent with our previous report (Takagi *et al*., 2019), in the presence of DCMU we previously did not detect a high residual component. The pH of the buffer used in the present experiments was pH 7.6, similar to *in vivo* stromal pH under light, whereas our previous experiment was conducted at pH 7.0, which is similar to the *in vivo* stromal pH in the dark (Trinh and Masuda, 2022). Therefore, we assessed the pH dependence of AA effects on the residual fluorescence in the presence of DCMU within a pH range of 6.8 to 8.0. The effect of AA on the residual fluorescence level was minimal under a lower pH, but strengthened with increase in the pH (**Fig. 2c**). The effect was greater at pH 8.0 than at pH 7.6, but the higher pH also slightly affected the Q_A_^−^ reoxidation kinetics in the presence of DCMU only. Therefore, we further studied the impacts of AA on PSII at pH 7.6.

### Photoinhibition of PSII by a single-turnover flash is observed under co-treatment with DCMU and AA

The residual, time-independent component of the Q_A_^−^ reoxidation kinetics has been ascribed to the equilibrium between Q_A_^−^ and Q_B_ in the absence of DCMU. However, in the presence of DCMU, DCMU binds to the Q_B_-site instead of PQ, and thus the residual component should reflect a different condition. Considering that the residual fluorescence level should correspond to the *F*_o_ level, we hypothesized that the significant increase in the residual component reflects photodamage to PSII induced by a single-turnover flash in the presence of DCMU and AA. Repetitive Q_A_^−^ reoxidation kinetics measurements were conducted with 5-min dark intervals, revealing that the residual component under co-treatment with DCMU and AA was accumulative and largely irreversible (**Extended Data Fig. 3**). Therefore, we conducted repetitive *F*_v_/*F*_m_ measurements of thylakoid membranes with 5-min dark intervals to study the photodamage to PSII induced by single-turnover saturating flashes (**Fig. 3a–c** and **Extended Data Fig. 4a**). In control and AA-treated thylakoid membranes in the absence of DCMU, *F*_v_/*F*_m_ remained almost unchanged, indicating that single-turnover flashes induced no notable photodamage. Only a slight decrease in *F*_v_/*F*_m_ was observed for thylakoid membranes incubated with only DCMU (after two flashes: *F*_v_/*F*_m_ > 0.7, 90% of initial *F*_v_/*F*_m_). On the other hand, under co-treatment with DCMU and AA, the *F*_v_/*F*_m_ of thylakoid membranes declined markedly after each single-turnover flash (after two flashes: *F*_v_/*F*_m_ = 0.53, 67% of initial *F*_v_/*F*_m_). This decline was mainly due to the increase in *F*_o_, as expected from the Q_A_^−^ reoxidation kinetics (**Fig. 3b, c**). Similar effects of DCMU and AA were observed when repetitive *F*_v_/*F*_m_ measurements were conducted with 40-min dark intervals (**Fig. 3a–c** and **Extended Data Fig. 4a**). The same effects of co-treatment with DCMU and AA were observed with isolated PSII membranes (**Fig. 3d–f** and **Extended Data Fig. 4b**). These results suggest that when both DCMU and AA are bound to PSII, PSII becomes extremely susceptible to photoinhibition, enabling even one single-turnover flash to severely damage PSII.

**Fig. 3.**
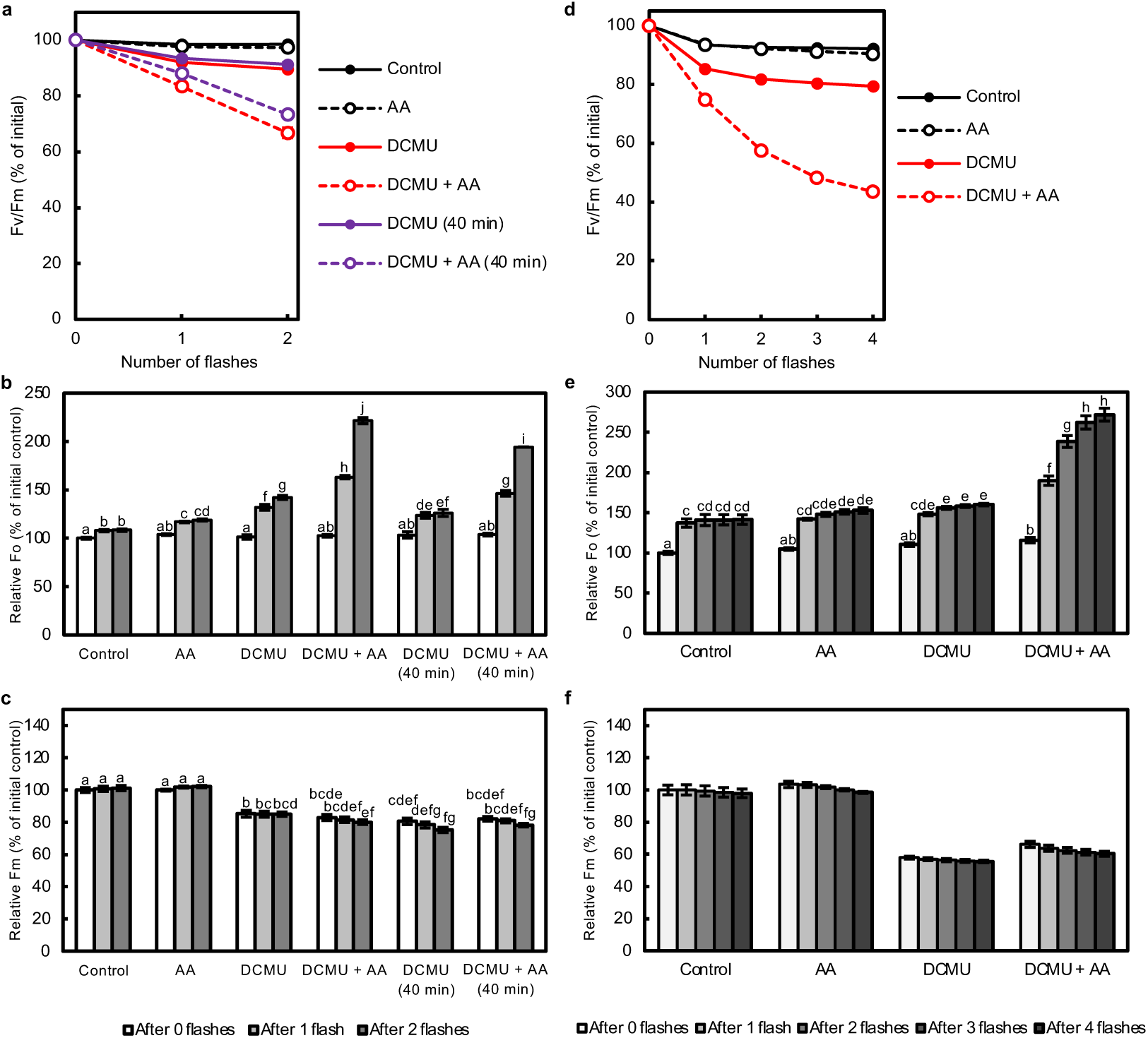
Photoinhibition of PSII induced by single-turnover flashes in the presence of DCMU and antimycin A (AA). Repetitive *F*_v_/*F*_m_ measurements as an indicator of PSII photoinhibition in (**a**–**c**) spinach thylakoid membranes and (**d**–**f**) isolated spinach PSII membranes treated with inhibitors. Change in the (**a**, **d**) *F*_v_/*F*_m_ values (% of initial value), (**b**, **e**) relative *F*_o_ values, and (**c**, **f**) relative *F*_m_ values after each single-turnover flash. (**a, d**) *F*_v_/*F*_m_ values of control samples (black filled circles) and samples treated with 10 µM AA (black open circles), 10 µM DCMU (red filled circles), 10 µM DCMU + 10 µM AA (red open circles) measured with 5-min dark intervals, and (**a**) 10 µM DCMU treated (purple filled circles) and 10 µM DCMU + 10 µM AA treated (purple open circles) samples measured with 40-min dark intervals. Data are mean ± SD (*n* = 3, technical replicates). Similar results were confirmed in independent experiments. In some instances, error bars are smaller than the symbol. (**b**, **c**, **e**) Different lowercase letters above bars indicate a statistically significant difference (*p* < 0.05, two-way ANOVA followed by Bonferroni–Holm’s test). (**f**) Two-way ANOVA showed no significant interaction effects, and only showed significant main effects of the herbicide type and number of flashes.

### AA1 and AA2 but not AA3 nor AA4 inhibit PSII

Commercially available AA is usually a mixture of closely related compounds, majorly antimycin A1, A2, A3, and A4 (**Fig. 4a**). We investigated whether the AA effects on PSII differ among the different AAs. Purified AA1–AA4 were used, as well as two commercially available AA mixtures isolated from *Streptomyces* sp. (AA mix#1 and AA mix#2) with separate lot numbers. In the absence of DCMU, Q_A_^−^ reoxidation was inhibited by AA1 and AA2, whereas little effect was observed with AA3 and AA4 (**Fig. 4b**). In the presence of DCMU, AA1 and AA2 strongly increased the residual fluorescence level, with AA1 showing a higher effect than AA2. In the meanwhile, AA3 and AA4 again had little effect on PSII (**Fig. 4c**). The effects of AA mix#1 and AA mix#2 on PSII were in between those of AA1 and AA3, with AA mix#1 which has a higher AA1 content having a stronger inhibitory effect than AA mix#2 does. The fact that slight structural differences among AAs lead to remarkably different inhibitory activities on PSII, implies a specific interaction between the AAs and PSII. Considering that among the four AAs, AA1 is the most hydrophobic and AA4 is the least, high hydrophobicity may also be important for the AA effects on PSII.

**Fig. 4.**
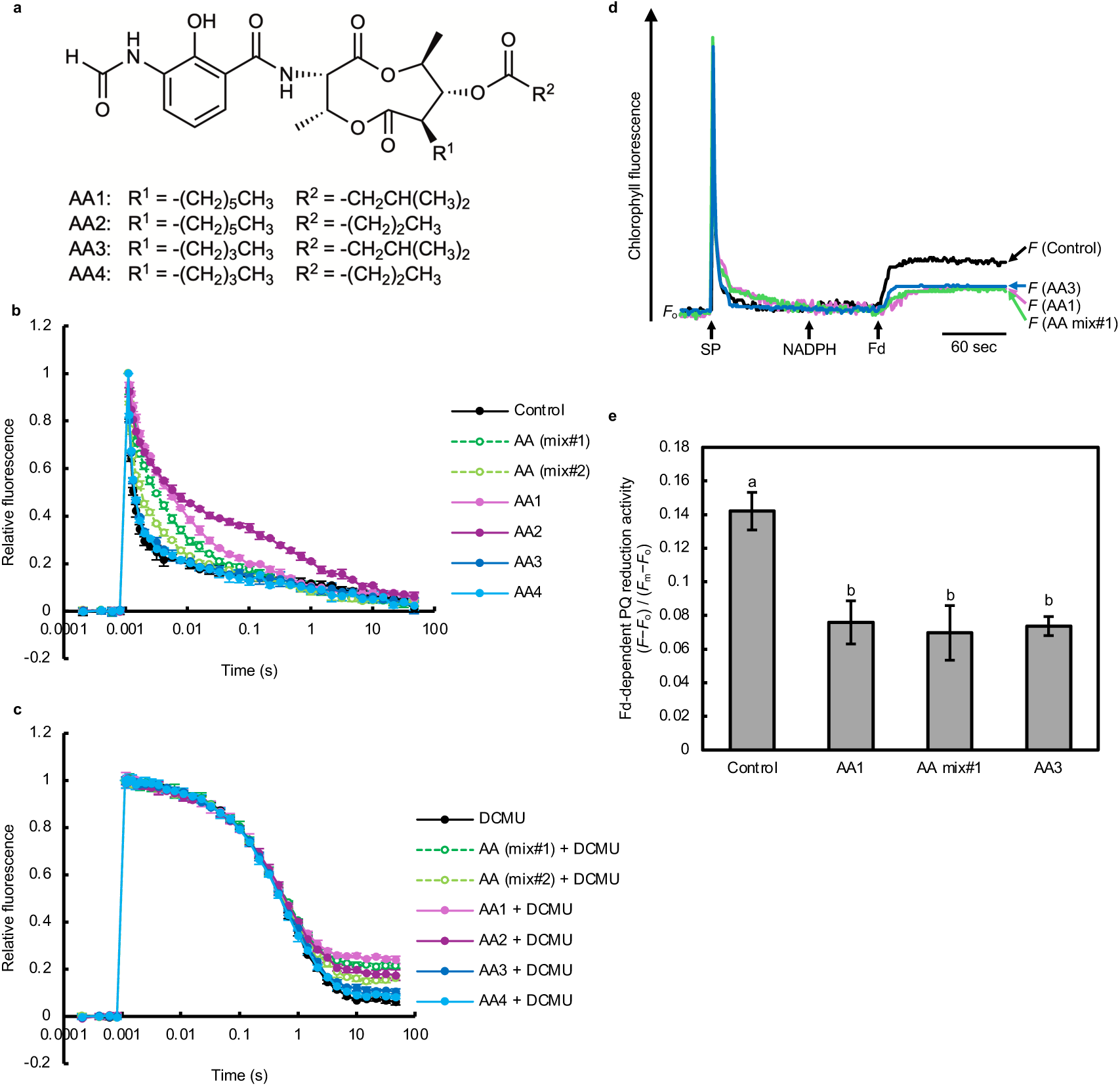
Effects of different antimycin A (AA) components on PSII and on Fd-dependent PQ reduction. (**a**) Structure of the major AA components: AA1, AA2, AA3, and AA4. (**b,c**) The effects of different AA components on the Q_A_^−^ reoxidation kinetics of PSII in isolated spinach thylakoid membranes. Control (black filled circles), 10 µM AA (two separate AA mixtures from *Streptomyces* sp.) (green and light green open circles), 10 µM AA1 (pink filled circles), 10 µM AA2 (purple filled circles), 10 µM AA3 (blue filled circles), and 10 µM AA4 (light blue filled circles) in the (**b**) absence and (**c**) presence of 10 µM DCMU. Data are mean ± SD (*n* = 3, technical replicates). Similar results were confirmed in independent experiments. (**d, e**) The effects of AA1, AA (mixture), and AA3 on Fd-dependent PQ reduction. (**d**) Increases in chlorophyll fluorescence by the addition of NADPH and Fd to ruptured chloroplasts without addition of AA (black line), or with addition of AA1 (pink line), AA mixture (green line), or AA3 Fd-dependent PQ reduction activity evaluated as (*F*−*F*_o_) / (*F*_m_−*F*_o_). Data are mean ± SD (*n* = 3, technical replicates). Similar results were confirmed in independent experiments. Different lowercase letters above bars indicate a statistically significant difference (*p* < 0.05, Tukey’s HSD test).

As AA is often used as an inhibitor of PGR5-dependent CEF-PSI, we also examined whether this pathway is sensitive to either or both of AA1 and AA3, by measuring the Fd-dependent PQ reduction activity in ruptured chloroplasts (**Fig. 4d, e**). AA1, AA mix#1, and AA3 all inhibited Fd-dependent PQ reduction to the same extent. A previous report also showed that AA3 inhibits PGR5-dependent CEF-PSI (Taira *et al*., 2013). This suggests that the inhibition of PGR5-dependent CEF-PSI by AA is likely to occur by a mechanism different from the inhibitory effect of AA on PSII. Moreover, this result shows that the use of AA3 instead of AA (mixtures) enables inhibition of PGR5-dependent CEF-PSI without notable effects on PSII.

### AA strongly affects PSII under co-treatment with DCMU, terbutryn, bromacil, or atrazine, but not with bromoxynil

DCMU is a urea-type inhibitor that binds to the Q_B_-site of PSII, but several other types of inhibitors bind at the Q_B_-site as well. To investigate whether the impact of AA on PSII can also be observed in the presence of Q_B_-site binding inhibitors other than DCMU, we examined the effects of AA on the Q_A_^−^ reoxidation kinetics and repetitive *F*_v_/*F*_m_ measurements in the presence of other Q_B_-site binding inhibitors: terbutryn (triazine-type), bromacil (uracil-type), and bromoxynil (phenol-type). Similar effects of AA on the Q_A_^−^ reoxidation kinetics were observed in the presence of terbutryn (10 µM) or bromacil (10 µM) instead of DCMU (10 µM). The residual fluorescence level was slightly higher under bromacil–AA co-treatment, and increased further by terbutryn–AA co-treatment, compared with DCMU–AA co-treatment (**Fig. 5a**). On the other hand, little effect of AA on the Q_A_^−^ reoxidation kinetics was observed in the presence of bromoxynil (10 µM). Co-treatment of AA with a higher concentration of bromoxynil (100 µM) similarly had no impact on the residual fluorescence. Consistent with this, the decrease in *F*_v_/*F*_m_ induced by single-turnover flashes was greatest with terbutryn–AA co-treatment, followed by bromacil–AA co-treatment and DCMU–AA co-treatment, whereas bromoxynil–AA co-treatment induced only a slight decrease in *F*_v_/*F*_m_ (**Fig. 5b** and **Extended Data Fig. 4c**). The decrease in *F*_v_/*F*_m_ under AA co-treatment with terbutryn, bromacil, or DCMU reflected a significant increase in *F*_o_, which was not observed with 100 µM bromoxynil + 10 µM AA treatment (**Fig. 5c, d**). The AA effects on the Q_A_^−^ reoxidation kinetics and repetitive *F*_v_/*F*_m_ measurements in the presence of atrazine were confirmed to be similar to those in the presence of DCMU (**Extended Data Fig. 5**). The impact of AA was much weaker, but still distinct, in the presence of bromoxynil instead of DCMU, including with isolated PSII membranes (**Extended Data Fig. 6**), suggesting that AA does bind to PSII together with bromoxynil. Bromoxynil, but not DCMU, downshifts the Q_A_^−^/Q_A_ redox potential (Krieger-Liszkay and Rutherford, 1998) and accelerates S_2_Q_A_^−^ recombination (Cser and Vass, 2007) (**Fig. 5a**). This may explain the difference in effects of AA in the presence of bromoxynil and the other Q_B_-site binding inhibitors.

**Fig. 5.**
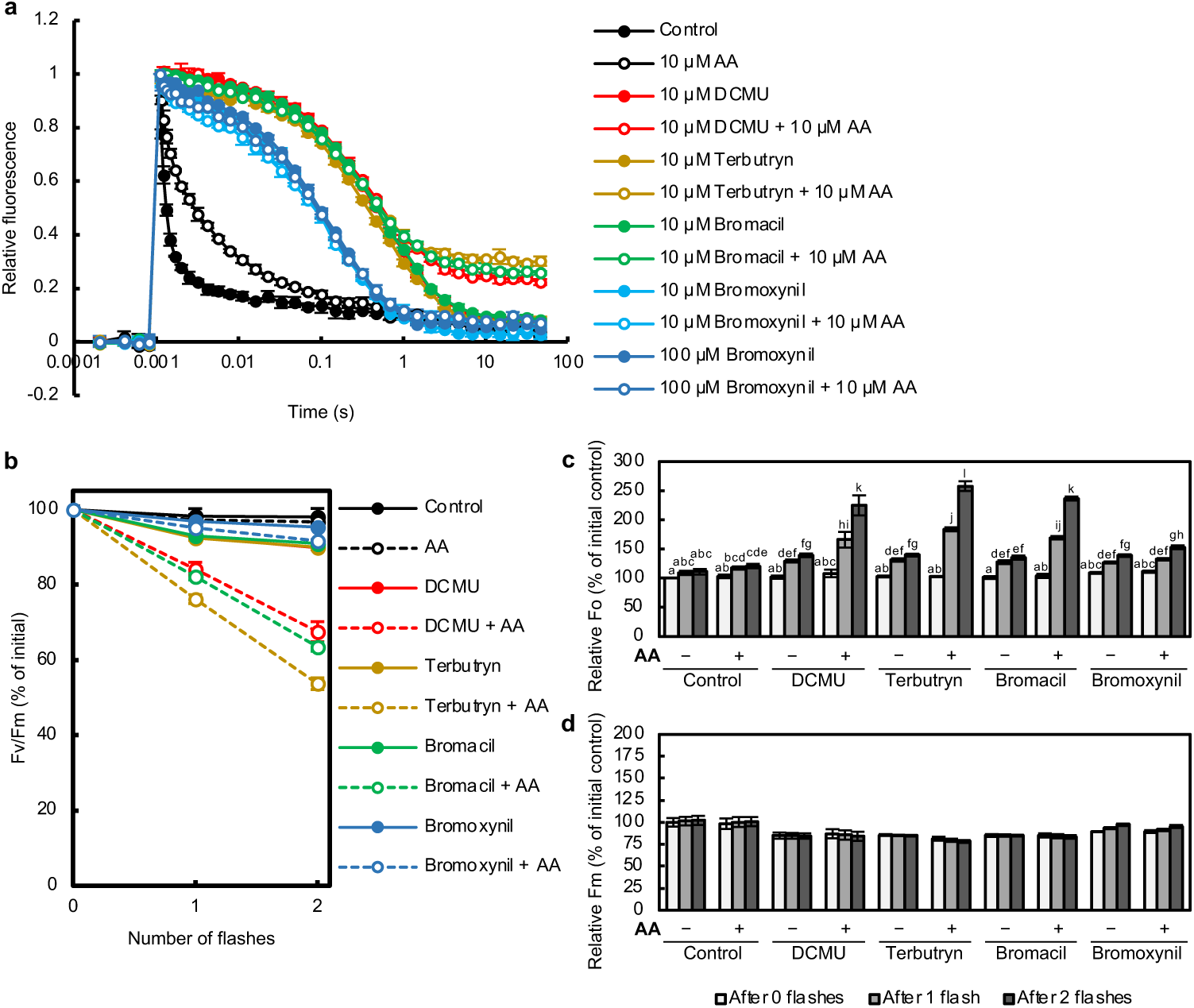
Effects of antimycin A (AA) on PSII in the presence of various Q_B_-site binding inhibitors. (**a**) Effects of AA on the Q_A_^−^ reoxidation kinetics of PSII in spinach thylakoid membranes in the absence or presence of DCMU, terbutryn, bromacil, or bromoxynil. (**b**–**d**) Repetitive *F*_v_/*F*_m_ measurements in spinach thylakoid membranes in the absence or presence of AA and Q_B_-site binding inhibitors. Change in the (**b**) *F*_v_/*F*_m_ values, (**c**) relative *F*_o_ values, and (**d**) relative *F*_m_ values after each single-turnover flash. (**a**) Q_A_^−^ reoxidation and (**b**) *F*_v_/*F*_m_ values (measured with 5-min dark intervals) of control samples (black filled circles), and samples treated with 10 µM AA (black open circles), 10 µM DCMU (red filled circles), 10 µM DCMU + 10 µM AA (red open circles), 10 µM terbutryn (yellow filled circles), 10 µM terbutryn + 10 µM AA (yellow open circles), 10 µM bromacil (green filled circles), 10 µM bromacil + 10 µM AA (green open circles), 100 µM bromoxynil (blue filled circles), 100 µM bromoxynil + 10 µM AA (blue open circles), and in **a**, 10 µM bromoxynil (light blue filled circles), 10 µM bromoxynil + 10 µM AA (light blue open circles). Data are mean ± SD (*n* = 3, technical replicates). Similar results were confirmed in independent experiments. In some instances, error bars are smaller than the symbol. (**c**) Different lowercase letters above bars indicate a statistically significant difference (*p* < 0.05, two-way ANOVA followed by Bonferroni–Holm’s test). (**d**) Two-way ANOVA showed no significant main effects of number of flashes or interaction

## Discussion

AA is frequently used as an inhibitor of the PGR5-dependent CEF-PSI in photosynthetic electron transport, although the mechanism is not known at all. In addition, AA affects the redox potential of Cyt *b*_559_ in PSII, which is considered to not participate in PGR5-dependent reactions. Nevertheless, such effects on PSII have been largely overlooked and AA is still often used as if it were a specific inhibitor of PGR5-dependent CEF-PSI in photosynthetic reactions. The present results have revealed that AA directly affects PSII and perturbs the electron transport within PSII. In particular, in the presence of Q_B_-site binding inhibitors DCMU, terbutryn, bromacil, or atrazine, AA strongly impacted on PSII, making PSII hypersensitive to light and enabling a single-turnover flash to induce severe photoinhibition. In addition, because Cyt *b*_559_ is considered to receive electrons from Q_B_, and the present results showed distinct, direct effects of AA on PSII in the presence of Q_B_-site binding inhibitors, the impacts of AA on PSII cannot be limited to Cyt *b*_559_.

The Q_A_^−^ reoxidation kinetics in the absence of DCMU suggested that AA suppresses electron transfer from Q_A_^−^ to Q_B_ and the binding of PQ to an empty Q_B_-site (**Fig. 2** and **Table 1**). This is evidently a direct effect of AA on PSII, as it was observed in thylakoid membranes (**Fig. 2a**) as well as in isolated PSII membranes (**Fig. 2b**). Together with our previous thermoluminescence measurements, which showed that AA affects the redox potential of Q_B_ rather than that of Q_A_ or donor side components (Takagi *et al*., 2019), we suggest that AA affects the environment surrounding the Q_B_-site, most likely by binding near the acceptor side of PSII. The Q_A_^−^ reoxidation measurements in the presence of very low concentration of DCMU, in which the effect of DCMU on PSII is not saturated, supports this suggestion because AA partially eliminated the effect of DCMU under this condition (**Extended Data Fig. 7**) although both DCMU and AA slowed Q_A_^−^ reoxidation (**Fig. 2**). This indicates that AA may affect the binding properties or binding affinity of DCMU to PSII. AA lowers the redox potential of Cyt *b*_559_, which is also located near the Q_B_-site (**Fig. 1**). Various mutations in Cyt *b*_559_ decrease the redox potential of Cyt *b*_559_ while affecting the redox properties of Q_B_ and/or delaying the electron transport from Q_A_^−^ to Q_B_ (Chiu *et al*., 2009, 2013; Hamilton *et al*., 2014; Nakamura, Boussac and Sugiura, 2019). Cyt *b*_559_ is positioned between two PQ exchange channels that connect the Q_B_-site with the outer side of PSII (Van Eerden *et al*., 2017). Inside or near these quinone exchange channels, additional quinone binding sites, Q_C_ and Q_D_, have been reported (Guskov *et al*., 2009; Kamada, Nakajima and Shen, 2023). These quinone binding sites are located close to the Q_B_-site and Cyt *b*_559_, and could affect their properties. Given that AA binds to the quinone binding site Q_i_ in complex III (Cyt *bc*_1_) in the mitochondrial respiratory chain, the Q_C_ and Q_D_ sites may be speculated as AA binding sites. This would be consistent with the strong effects of AA1 and AA2 which have longer hydrophobic tails mimicking PQ, in contrary to AA3 and AA4 with shorter tails having little effect on PSII (**Fig. 4a–c**). Moreover, we found that myxothiazol, a well-known inhibitor that binds to the Q_o_ quinone binding site of the mitochondrial complex III, also inhibits PSII in a similar way as AA does in the absence of DCMU (**Extended Data Fig. 8**). This further supports that AA (and myxothiazol) may bind to PQ binding sites in PSII other than the Q_A_- and Q_B_-sites. The mechanism by which AA suppresses reactions in PSII in the absence of Q_B_-site binding inhibitors may be a mixed result of perturbation of the protein environment around the Q_B_-site, blockage of the PQ exchange channels, and inhibition of electron transfer pathways within PSII such as cyclic electron flow within PSII mediated by Cyt *b*_559_ (Takagi *et al*., 2019) and/or Q_C_ (Gates *et al*., 2022).

Repetitive *F*_v_/*F*_m_ measurements revealed an intriguing effect of AA on PSII in the presence of DCMU (**Fig. 3**). The presence of DCMU only or AA only had little effect on *F*_v_/*F*_m_, whereas co-treatment with DCMU and AA caused a marked decline in *F*_v_/*F*_m_ after every flash with 5- or 40-min dark intervals, implying severe photodamage to PSII. This decline in *F*_v_/*F*_m_ was due to a remarkable increase in *F*_o_, consistent with the high residual fluorescence in Q_A_^−^ reoxidation measurements (**Fig. 3b, c, e, f**). *F*_o_ increased after every flash illumination and little recovery was observed even after 40-min incubation in the dark. The rise in *F*_o_ after each flash can be due to the inactivation of PSII reaction centers (Yamane *et al*., 1997; Hong and Xu, 1999; Yi *et al*., 2006). This may also be partially attributed to a stable Q_A_^−^ formed by P680^+^ reduction by electron donors (Mn^2+^, TyrD, or Cyt *b*_559_/Chl_Z_/Car_D2_) in PSII that lost a functional Mn_4_CaO_5_ cluster (Nixon, Trost and Diner, 1992; Debus *et al*., 2000; Viola *et al*., 2022) or in which the Mn_4_CaO_5_ cluster is reduced from S_2_ to S_1_ owing to an ADRY (acceleration of the deactivation reactions of the water-splitting enzyme system Y)-like effect (Yerkes and Crofts, 1992). However, while even the very slow TyrD^+^Q_A_^−^ recombination has been reported to occur with a half-time of ∼10 min (Johnson, Boussac and Rutherford, 1994), the *F*_o_ level remained high even after 40 min (**Fig. 3b**). Such extremely stable Q_A_^−^ has only been characterized after PSII photoinhibition by multiple turnovers under anaerobic conditions (Vass *et al*., 1992), which is a completely different case. Therefore, although the slightly lower *F*_o_ with 40-min intervals compared with those with 5-min intervals may reflect oxidation of stable Q_A_^−^ by slow charge recombination, the high *F*_o_ level with 40-min intervals more likely results from irreversibly photodamaged PSII. This photodamage was induced by AA in the presence of DCMU, terbutryn, bromacil, or atrazine, but not in the presence of bromoxynil (**Fig. 5** and **Extended Data Fig. 5**), suggesting it can be avoided by a rapid S_2_Q_A_^−^ recombination by downshifting the redox potential of Q_A_^−^/Q_A_ (Krieger-Liszkay and Rutherford, 1998). This is consistent with the pH dependence of the AA effects (**Fig. 2c**), because within the pH range of around 7 to 8, faster S_2_Q_A_^−^ recombination at lower pH has been reported (Robinson and Crofts, 1984; Vass and Inoue, 1986). While further spectroscopic and structural studies are required to elucidate the detailed mechanism of the effect of AA in the presence of DCMU, it is apparent that a large proportion of PSII becomes nonfunctional even by a single flash illumination under co-treatment with DCMU and AA.

The impacts of AA on the Q_A_^−^ reoxidation kinetics of isolated PSII membranes (**Fig. 2b**), the decrease in *F*_v_/*F*_m_ of PSII membranes after each single-turnover flash in the presence of DCMU and AA (**Fig. 3d**), and the difference in AA effects on PSII with different Q_B_-site binding inhibitors (**Fig. 5** and **Extended Data Fig. 5**) demonstrate that AA has direct, non-negligible impacts on PSII. The effects of inhibition of PGR5-dependent CEF-PSI using AA are often assayed by measuring chlorophyll fluorescence, which largely derives from PSII. In addition, AA is occasionally used in conjunction with DCMU to investigate the effects of PGR5-dependent CEF-PSI while the electron flow from PSII to the PQ pool is blocked. The present results imply that the effects of AA on photosynthesis are not attributable simply to inhibition of PGR5-dependent CEF-PSI. In fact, several reports have noted that AA influences PSII in *pgr5* mutants (Havaux, Rumeau and Ducruet, 2005; Takahashi *et al*., 2009; Kono and Terashima, 2016). Undoubtedly considering the consequences of AA treatment as a result of PGR5-dependent CEF-PSI inhibition may lead to crucial misinterpretation. Re-interpretation of previous results obtained with AA, considering its effects on PSII, may help improve our understanding of PGR5 functions.

While our findings suggest that the use of AA as an inhibitor of PGR5-dependent CEF-PSI is inadvisable, we propose that the use of AA3 is useful for distinguishing the effects of AA on PSII and on PGR5-dependent CEF-PSI. Q_A_^−^ reoxidation measurements with different components of AA revealed that of the four major AA components, only AA1 and AA2 possess notable PSII inhibiting activity (**Fig. 4b, c**). Compared to AA1, AA2, or AA (mixtures), the two AA components, AA3 and AA4, showed very little effect on PSII. Meanwhile, AA3 inhibited PGR5-dependent CEF-PSI to the same rate as AA1 and AA (mixtures) do (**Fig. 4d, e**). Therefore, using AA3 instead of AA (mixtures) would allow observation of the actual consequences of inhibition of PGR5-dependent CEF-PSI without affecting PSII.

AA3, as well as AA mixtures, also inhibits the mitochondrial complex III (Dickie *et al*., 1963; Miyoshi *et al*., 1991; Tokutake *et al*., 1994, p. 19). Currently, inhibition of complex III is usually conducted by using either AA (mixtures) or myxothiazol. As AA mixtures have a strong effect on both PSII and PGR5-dependent CEF-PSI, the use of myxothiazol would be preferable, although myxothiazol is not completely specific to complex III either (**Extended Data Fig. 8**). Our results suggest that combining the results obtained by myxothiazol treatment and by AA3 treatment can be effective for distinguishing the effects of complex III inhibition, PGR5-dependent CEF-PSI inhibition, and PSII inhibition.

## Materials and methods

### Isolation of PSII membranes and thylakoid membranes

Oxygen-evolving PSII membranes (BBY membranes) (Berthold, Babcock and Yocum, 1981) were isolated from market spinach using a previously reported method (Imaizumi *et al*., 2022) following Yamamoto et al. (Yamamoto, Leng and Shen, 2011). Spinach thylakoid membranes were isolated from market spinach by modifying the same method (Yamamoto, Leng and Shen, 2011; Imaizumi *et al*., 2022). *Chlamydomonas reinhardtii* thylakoid membranes were isolated from the wild-type *C. reinhardtii* strain CC-125 based on a previously reported method (Nishimura, Sato and Ifuku, 2017). *Chlamydomonas* cells were grown in liquid tris-acetate-phosphate (TAP) medium at 25 °C with continuous light (30 µmol photons m^−2^ s^−1^), and the cells were harvested by centrifugation at 1000×*g*. After resuspending the cells in buffer C1 (25 mM HEPES-NaOH, 0.4 M sucrose, 30 mM NaCl, 5 mM MgCl_2_, pH7.0), the cells were disrupted with glass beads (100 µm diameter) using a Mini-Beadbeater (BioSpec Products). Thylakoid membranes were obtained by centrifugation at 20,400×*g*, and were suspended in buffer C2 (25 mM HEPES-NaOH, 20 mM NaCl, 12.5% (v/v) glycerol, pH 7.0).

### Isolation of intact chloroplasts

Spinach leaves were homogenized in ice-cold grinding buffer (50 mM HEPES-KOH, 330 mM sorbitol, 1 mM MnCl_2_, 1 mM MgCl_2_, 2 mM EDTA, 0.03% (w/v) bovine serum albumin, pH 8.0). The homogenate was filtered through four layers of Miracloth, and centrifuged at 2300×*g* for 1 min. The pellet was resuspended in grinding buffer, layered onto a discontinuous 40% / 80% Percoll density gradient, and centrifuged at 2100×*g* for 20 min. Intact chloroplasts, which formed a band at the interface between 40 and 80% Percoll, were collected and washed in resuspension buffer (50 mM HEPES-KOH, 330 mM sorbitol, 1 mM MnCl_2_, 1 mM MgCl_2_, 2 mM EDTA, pH 8.0).

### Inhibitors

AA (mixture of AAs isolated from *Streptomyces* sp.) was purchased from Sigma-Aldrich (product number: A8674). According to the Certificate of Analysis by Sigma-Aldrich, the major components of AA mix#1 (lot number: 125K4030) were AA1 (73%), AA2 (19%), AA3 (5%), and AA4 (1%), and those of AA mix#2 (batch number: 0000187236) were AA1 (34%), AA2 (17%), AA3 (34%), and AA4 (13%). AA mix#1 was used as AA unless indicated. AA1–AA4 were purchased from BioAustralis Fine Chemicals, and the product numbers and batch numbers were as follows: AA1 (BIA-A1442, batch AC33.74), AA2 (BIA-A1443, batch HC11.47), AA3 (BIA-A1444, batch AC32.10), AA4 (BIA-A1445, batch AC32.78). All inhibitor stock solutions were prepared in ethanol.

### Measurement of Q_A_^−^ reoxidation kinetics

The increase and subsequent decay of chlorophyll fluorescence induced by a single-turnover flash was monitored using a double-modulation fluorometer FL3500 (Photon Systems Instruments) based on methods reported previously (Reifarth *et al*., 1997; Vass, Kirilovsky and Etienne, 1999). Prior to measurements, thylakoid membranes or PSII membranes (5 µg Chl mL^−1^ unless indicated) were incubated with inhibitor compounds (e.g., DCMU and AA) in buffer A (20 mM HEPES-NaOH, 1 M betaine, 10 mM NaCl, 5 mM MgCl_2_, pH 7.6 (unless indicated)) for 3 min at room temperature in complete darkness. Data were fit to a tri-exponential decay equation after normalization (Reifarth *et al*., 1997) using the software OriginPro.

### Repetitive measurement of *F*_v_/*F*_m_

Dark-adapted thylakoid membranes or PSII membranes (10 µg Chl mL^−1^ unless indicated) were incubated with inhibitor compounds in buffer A for 2 min at room temperature in complete darkness. *F*_v_/*F*_m_ was measured using AquaPen-C AP-C 100 fluorometer (Photon Systems Instruments). Between each *F*_v_/*F*_m_ measurement, samples were kept in complete darkness for 5 min (unless indicated).

### *In vitro* Fd-dependent PQ reduction assay

The *in vitro* Fd-dependent PQ reduction assay was performed based on previous reports (Endo *et al*., 1998; Okegawa and Motohashi, 2020). Freshly prepared intact chloroplasts (20 µg Chl mL^−1^) were osmotically ruptured in rupture buffer (50 mM HEPES-KOH, 7 mM MgCl_2_, 1 mM MnCl_2_, 2 mM EDTA, 30 mM KCl, 0.25 mM KH_2_PO_4_, pH 7.6) just before measurement. AA1, AA mix#1, or AA3 were added to a final concentration of 10 µM, leading to a final concentration of 0.2% (v/v) ethanol. 250 µM NADPH (Oriental Yeast) and 5 µM spinach Fd (Sigma-Aldrich) were used as electron donors. The chlorophyll fluorescence levels under weak light (0.9 µmol photons m^−2^ s^−1^) were monitored by the Photosynthesis Yield Analyzer MINI-PAM (Walz), and the activity of Fd-dependent PQ reduction was assayed by measuring the increases in chlorophyll fluorescence by addition of NADPH and Fd. The fluorescence levels were normalized by the *F*_m_ levels and standardized by the *F*_o_ levels, and the Fd-dependent PQ reduction activity was evaluated using the equation (*F*−*F*_o_)/(*F*_m_−*F*_o_) (Okegawa and Motohashi, 2020) where *F* is the fluorescence level 2 min after addition of both NADPH and Fd.

## Acknowledgements

This work was supported in part by JSPS Grant-in-Aid for JSPS Fellows JP23KJ1361 to K. Imaizumi and for Challenging Research (Exploratory) JP24K21968 to K. Ifuku. We thank Robert McKenZie, from Edanz (https://jp.edanz.com/ac) for editing a draft of this manuscript.

## Author contributions

K. Imaizumi, D. Takagi, and K. Ifuku conceived the project; K. Imaizumi performed all analyses and drafted the original manuscript; K. Imaizumi, and K. Ifuku revised the manuscript and wrote the final manuscript, and all authors joined the discussion of the results.

---

**Extended Data Fig. 1.**
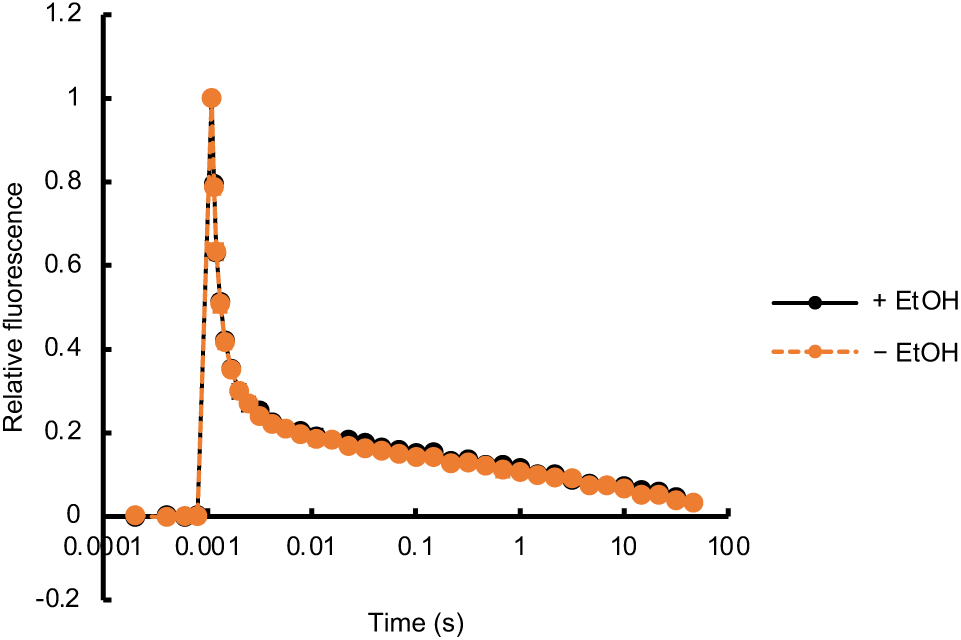
Effects of 0.2% ethanol on the Q_A_^−^ reoxidation kinetics of PSII. Q_A_^−^ reoxidation kinetics measurements of spinach thylakoid membranes were conducted with (black circles) and without (orange circles) 0.2% (v/v) ethanol. Data are mean ± SD (*n* = 3, independent experiments). In some instances, error bars are smaller than the symbol.

**Extended Data Fig. 2.**
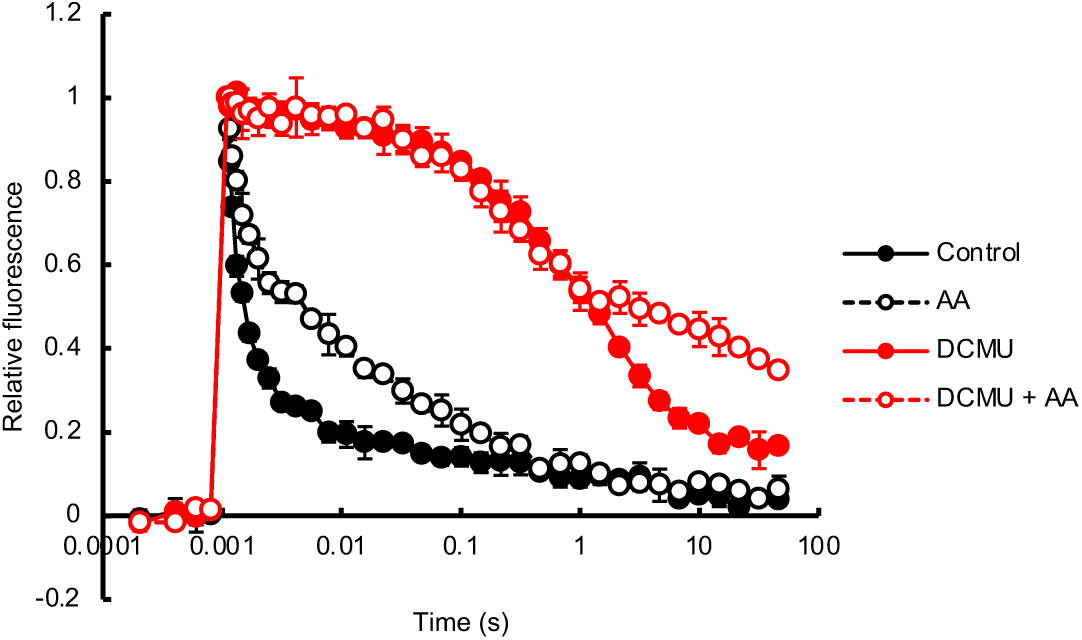
Effects of antimycin A (AA) on the Q_A_^−^ reoxidation kinetics of PSII using thylakoid membranes isolated from *Chlamydomonas reinhardtii*. Effects of AA on PSII in *C. reinhardtii* thylakoid membranes in the absence or presence of DCMU. Control (black filled circles), 10 µM AA (black open circles), 10 µM DCMU (red open circles), 10 µM DCMU + 10 µM AA (red open circles). Data are mean ± SD (*n* = 3, technical replicates). Similar results were confirmed in independent experiments. In some instances, error bars are smaller than the symbol.

**Extended Data Fig. 3.**
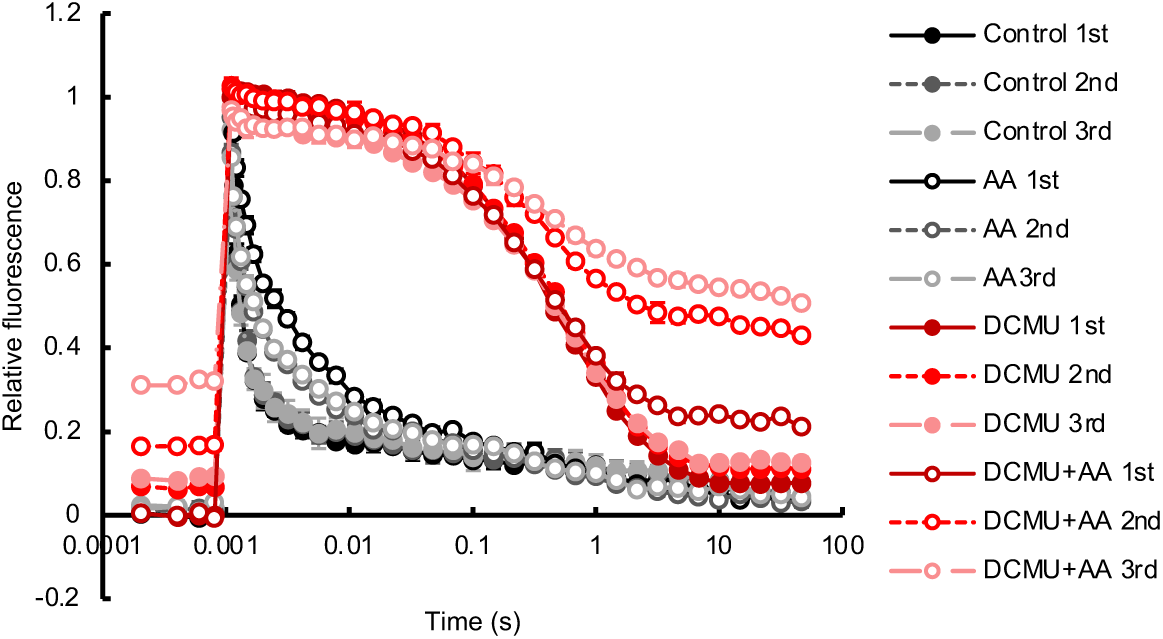
Repetitive measurement of the Q_A_^−^ reoxidation kinetics of PSII. Repetitive Q_A_^−^ reoxidation kinetics measurements of PSII in spinach thylakoid membranes were conducted with 5-min dark intervals in the absence (black, gray, and light gray) or presence (dark red, red, and pink) of 10 µM DCMU with (open circles) or without (filled circles) 10 µM AA. Data are mean ± SD (*n* = 3, technical replicates). Similar results were confirmed in independent experiments. In some instances, error bars are smaller than the symbol.

**Extended Data Fig. 4.**
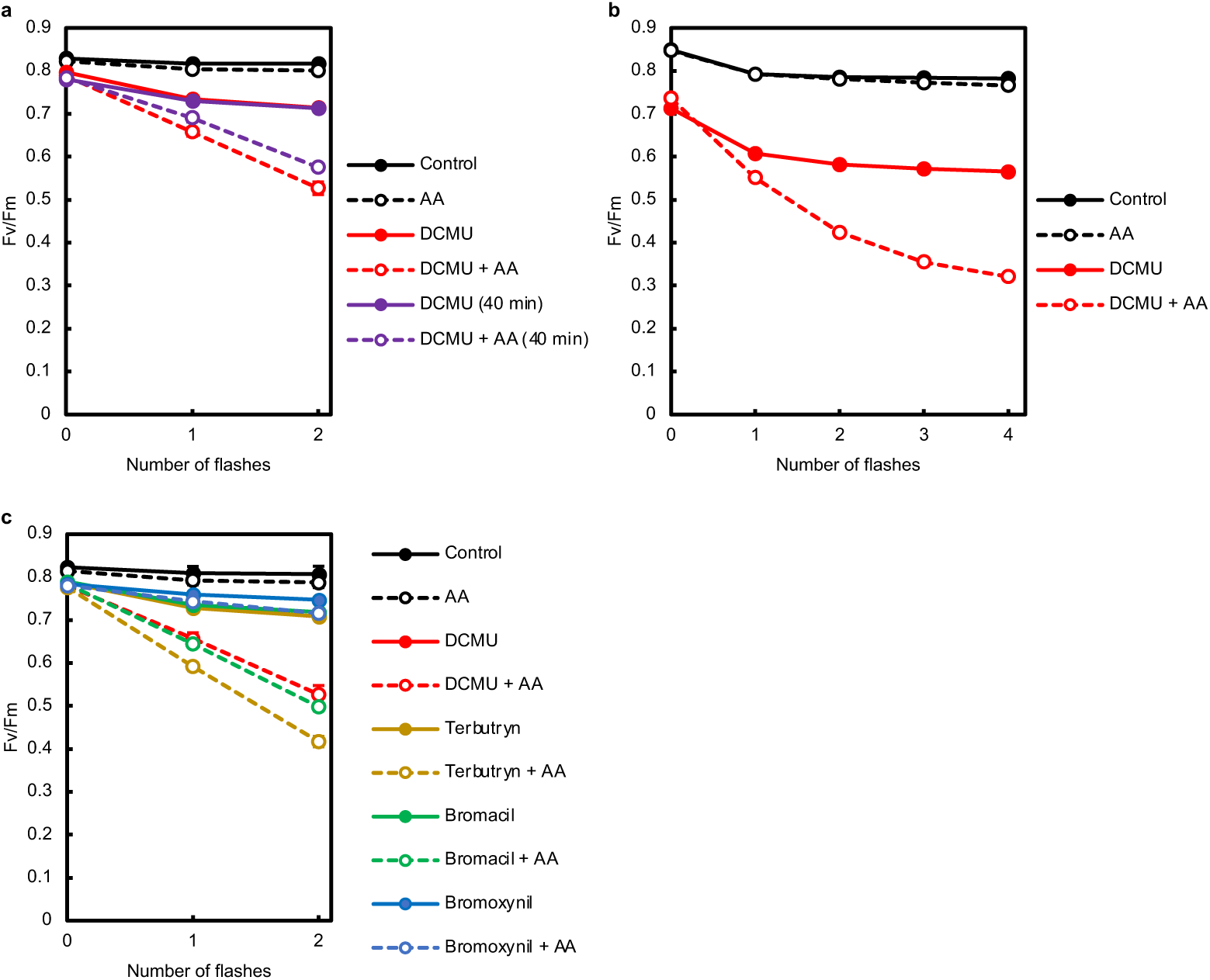
Actual *F*_v_/*F*_m_ values in repetitive *F*_v_/*F*_m_ measurements. Actual *F*_v_/*F*_m_ values corresponding to (**a**) Figure 3a, (**b**) Figure 3d, and (**c**) Figure 4c. Data are mean ± SD (*n* = 3, technical replicates). Similar results have been confirmed in independent experiments. In some instances, error bars are smaller than the symbol.

**Extended Data Fig. 5.**
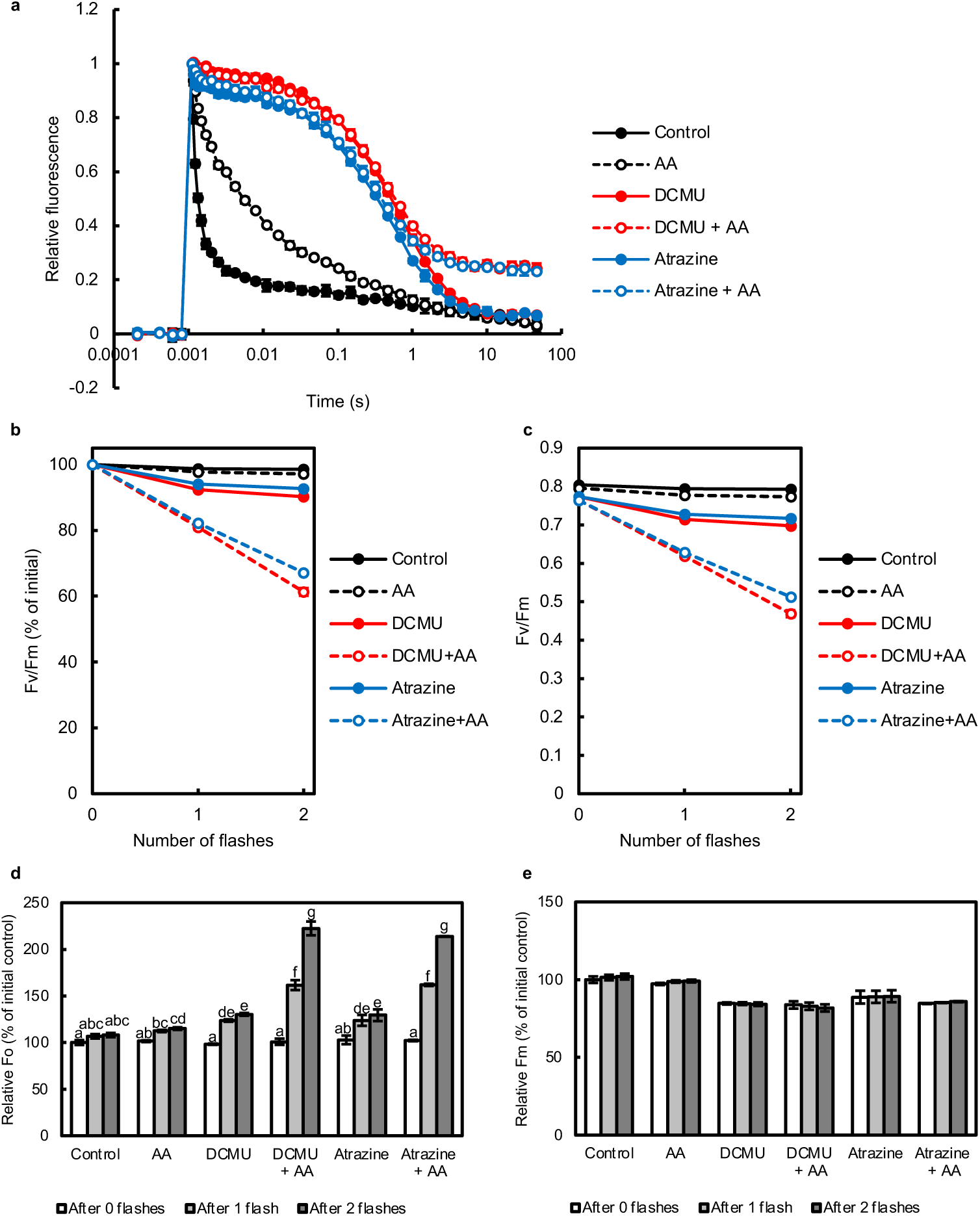
Effects of antimycin A (AA) on PSII in the presence of atrazine. (**a**) Effects of AA on the Q_A_^−^ reoxidation kinetics of PSII in spinach thylakoid membranes in the absence or presence of DCMU or atrazine. (**b–e**) Repetitive *F*_v_/*F*_m_ measurements in spinach thylakoid membranes in the absence or presence of AA, DCMU, or atrazine. Change in the (**b**, **c**) *F*_v_/*F*_m_ values (**b**, % of initial; **c**, actual value), (**d**) relative *F*_o_ values, and (**e**) relative *F*_m_ values after each single-turnover flash with 5-min dark intervals. (**a**) Q_A_^−^ reoxidation and (**b**, **c**) *F*_v_/*F*_m_ values of control samples (black filled circles), and samples treated with 10 µM AA (open black circles), 10 µM DCMU (red filled circles), 10 µM DCMU + 10 µM AA (red open circles), 10 µM atrazine (blue filled circles), and 10 µM atrazine + 10 µM AA (blue open circles). Data are mean ± SD (*n* = 3, technical replicates). Similar results were confirmed in independent experiments. In some instances, error bars are smaller than the symbol. (**d**) Different lowercase letters above bars indicate a statistically significant difference (*p* < 0.05, two-way ANOVA followed by Bonferroni– Holm’s test). (**e**) Two-way ANOVA showed no significant main effects of number of flashes or interaction effects, and only showed significant main effects of the herbicide type.

**Extended Data Fig. 6.**
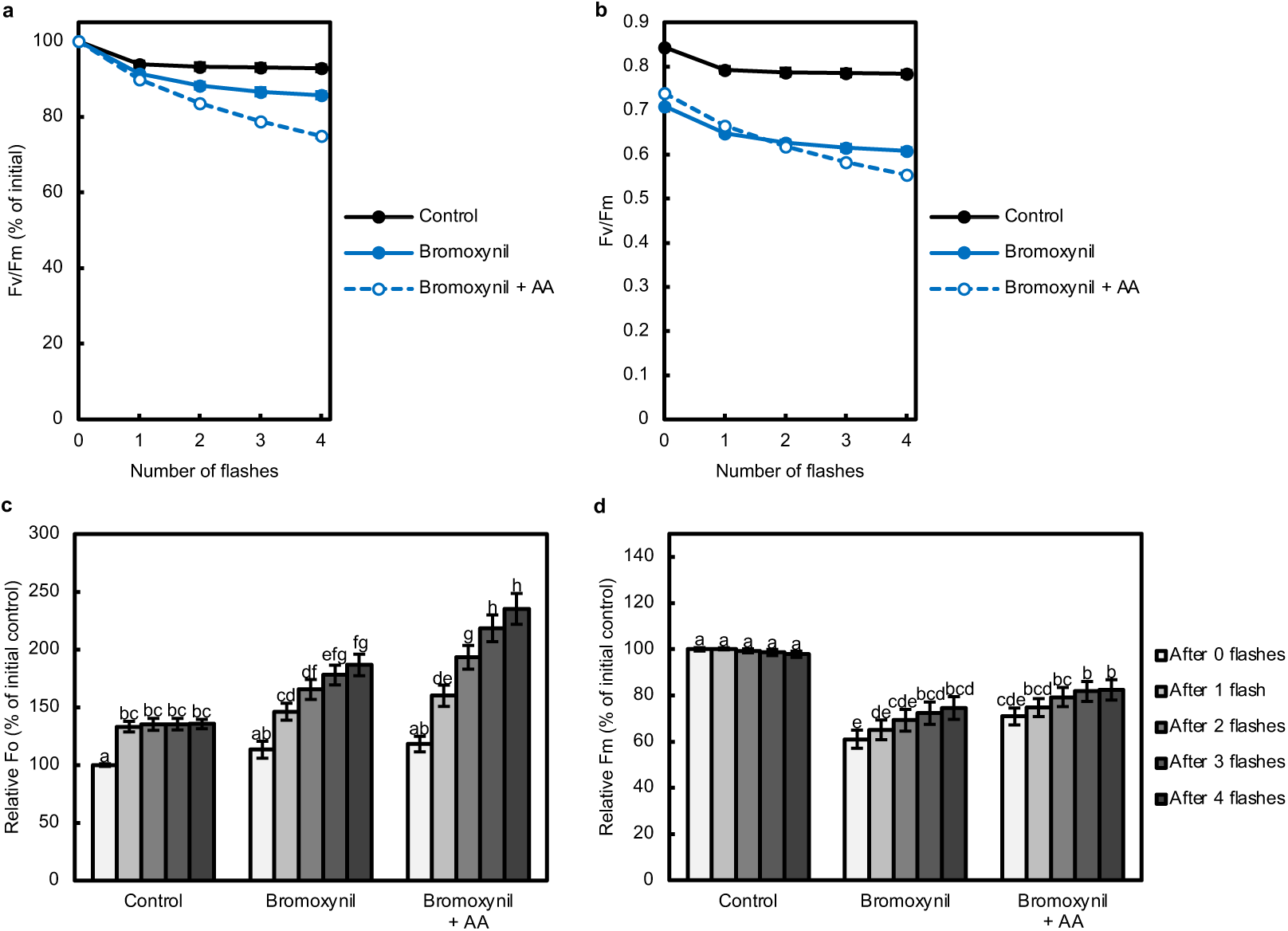
Effects of antimycin A (AA) on PSII in the presence of bromoxynil. The effects of AA on repetitive *F*_v_/*F*_m_ measurements of isolated PSII membranes were assayed in the presence of bromoxynil. Change in the (**a**, **b**) *F*_v_/*F*_m_ values (**a**, % of initial; **b**, actual value), (**c**) relative *F*_o_ values, and (**d**) relative *F*_m_ values after each single-turnover flash with 5-min dark intervals. (**a**, **b**) *F*_v_/*F*_m_ of control samples (black filled circles) and samples treated with 100 µM bromoxynil (blue filled circles), and 100 µM bromoxynil + 10 µM AA (blue open circles). (**c**, **d**) Different lowercase letters indicate a statistically significant difference (*p* < 0.05, two-way ANOVA followed by Bonferroni-Holm’s test). In all panels, data are mean ± SD (*n* = 3, technical replicates). Similar results were confirmed in independent experiments. In some instances, error bars are smaller than the symbol.

**Extended Data Fig. 7.**
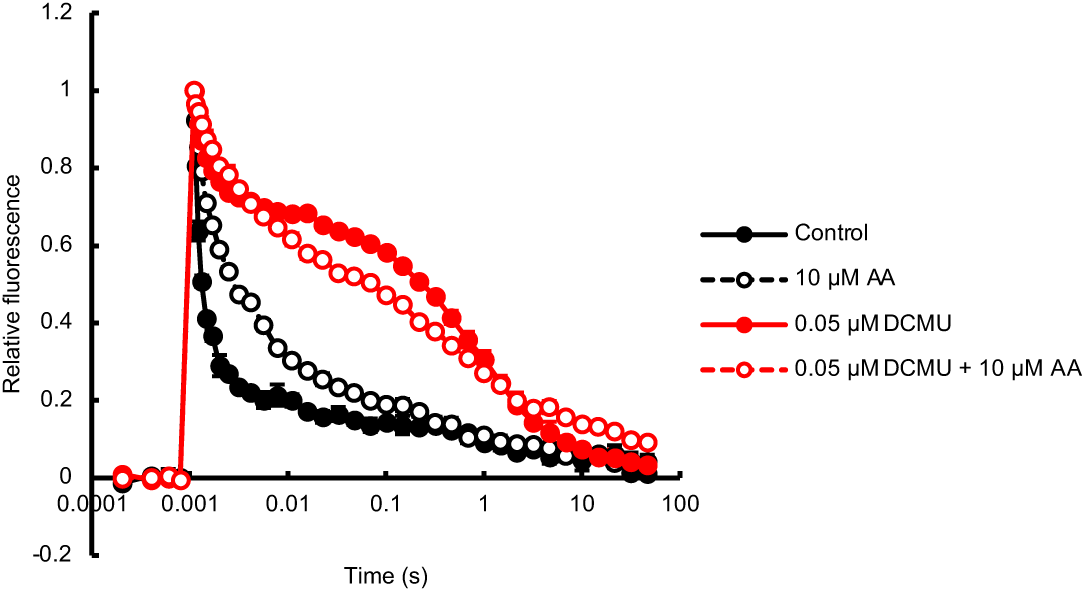
Effects of antimycin A (AA) on the Q_A_^−^ reoxidation kinetics of PSII under a low concentration of DCMU. Effects of AA on PSII in spinach thylakoid membranes in the absence or presence of DCMU at a low concentration in which the effects of DCMU are non-saturated. Control (black filled circles), 10 µM AA (black open circles), 0.05 µM DCMU (red open circles), 0.05 µM DCMU + 10 µM AA (red open circles). Data are mean ± SD (*n* = 3, technical replicates). Similar results were confirmed in independent experiments. In some instances, error bars are smaller than the symbols.

**Extended Data Fig. 8.**
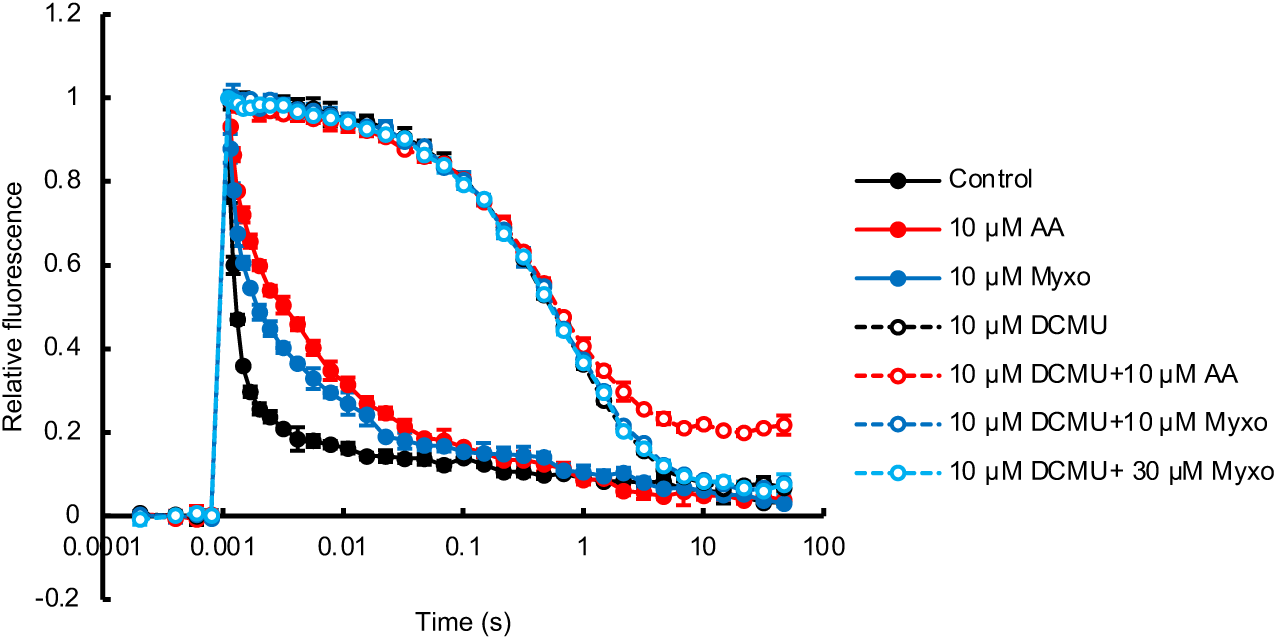
Effects of antimycin A (AA) and myxothiazol (Myxo) on the Q_A_^−^ reoxidation kinetics of PSII. Effects of AA and Myxo on PSII in thylakoid membranes in the absence or presence of DCMU. Control (black filled circles), 10 µM AA (red filled circles), 10 µM Myxo (blue filled circles), 10 µM DCMU (black open circles), 10 µM DCMU + 10 µM AA (red open circles), 10 µM DCMU + 10 µM Myxo (blue open circles), 10 µM DCMU + 30 µM Myxo (light blue open circles). Data are mean ± SD (*n* = 3, technical replicates). Similar results were confirmed in independent experiments. In some instances, error bars are smaller than the symbols.

